# Using QC-Blind for quality control and contamination screening of bacteria DNA sequencing data without reference genome

**DOI:** 10.1101/438655

**Authors:** Wang Xi, Yan Gao, Zhangyu Cheng, Chaoyun Chen, Maozhen Han, Kang Ning

**Author notes:** These authors contributed equally to this work (Wang Xi, Yan Gao).

## Abstract

Quality control in next generation sequencing has become increasingly important as the technique becomes widely used. Tools have been developed for filtering possible contaminants in the sequencing data of species with known reference genome. Unfortunately, reference genomes for all the species involved, including the contaminants, are required for these tools to work. This precludes many real-life samples that have no information about the complete genome of the target species, and are contaminated with unknown microbial species.

In this work we propose QC-Blind, a novel quality control pipeline for removing contaminants without any use of reference genomes. The pipeline requires only very little information from the marker genes of the target species. The entire pipeline consists of unsupervised read assembly, contig binning, read clustering and marker gene assignment.

When evaluated on *in silico*, *ab initio* and *in vivo* datasets, QC-Blind proved effective in removing unknown contaminants with high specificity and accuracy, while preserving most of the genomic information of the target bacterial species. Therefore, QC-Blind could serve well in situations where limited information is available for both target and contamination species.

**IMPORTANCE:** At present, many sequencing projects are still performed on potentially contaminated samples, which bring into question their accuracies. However, current reference-based quality control method are limited as they need either the genome of target species or contaminations. In this work we propose QC-Blind, a novel quality control pipeline for removing contaminants without any use of reference genomes. When evaluated on *in silico*, *ab initio* and *in vivo* datasets, QC-Blind proved effective in removing unknown contaminants with high specificity and accuracy, while preserving most of the genomic information of the target bacterial species. Therefore, QC-Blind is suitable for real-life samples where limited information is available for both target and contamination species.

## INTRODUCTION

As next generation sequencing (NGS) techniques, such as those based on the Illumina platform[1], become more widely used, the need for accuracy has likewise become increasingly urgent. At present, many sequencing projects are still performed on potentially contaminated samples, which bring into question their accuracies.

In most research scenarios, a target species is considered as the organism under study, while other species are seen as contaminants. These contaminants often include the microbial species found in the environment of the molecular biology laboratories[2–4]. Their interference with the sequencing would compromise the precision and reproducibility of the analysis[2]. Usually, these contaminations consist of not one, but a mixture of microbial species[5]. Previous studies in the removal of contaminants are concerned with water or soil samples, where relatively comprehensive bacteria genera profiles exist[1, 2]. However, contaminants removal becomes difficult when the profile of the sample is unknown[6, 7]. The problem becomes even more difficult when the reference genomes for both the target and the contaminations are not available[5, 8].

One strategy to identify and remove contaminants is the metagenomic approach[9], which facilitate taxonomical and functional analyses of the contaminating microbial genomes. A few methods based on this strategy have been proposed: SourceTracker[10] applies Bayesian inference to estimate the composition and abundance of microbial contaminations, while DeconSeq[11] deals with possible contamination from human through long reads alignment; and QC-Chain. We have also published a method to differentiate the reads from target species and contaminations[12], based on contig clustering[13]. However, the false positive rate of read assignment remained high, and potentially valuable information were not considered. For instance, knowledge of the abundance correlation of a certain target species among multiple samples (with similar contaminations) were not utilized[12].

In this study, we propose QC-Blind, a pipeline for bacteria NGS data quality control and contamination screening with high specificity and accuracy. The pipeline not only reduces the false positive rate for read assignment, but also requires only a few marker genes for differentiating reads from target bacteria species and bacteria contaminations. QC-Blind could remove contaminations that were introduced during sample handling, and recover genomes from mixed cultures / environmental samples. Extensive downstream performance evaluations based on *in silico*, *ab initio* and *in vivo* datasets, showed the method to be effective. As most microbial contaminations could be removed and almost complete genomic information of target bacteria species could be preserved after processing, this pipeline has shown near optimal solution for quality control and contamination screening of bacteria DNA sequencing data.

## MATERIALS AND METHODS

The general process flow of QC-Blind is as follows. First, reads are assembled into contigs. The contigs are clustered into species-level groups by species abundance and sequence features. Then, the marker genes of the target species (generated through MetaPhlAn2[14] and manual curation) are utilized to identify the contig clusters for the target species. Here, the main issues to sort out are with regard to the assembly and clustering accuracies, as well as the specificity of the contig clusters for target species. To perform this fine-tuning, we put the method to very thorough test on simulated, *ab initio* and *in vivo* datasets. The simulated and sequencing data are deposited to NCBI SRA with project access number PRJNA491366.

### 1. Simulated and Real Datasets

Three types of metagenomic datasets have been utilized in this study: simulated, *ab initio* and *in vivo* (**Figure 1**, **Table 1**).

**Table 1.**
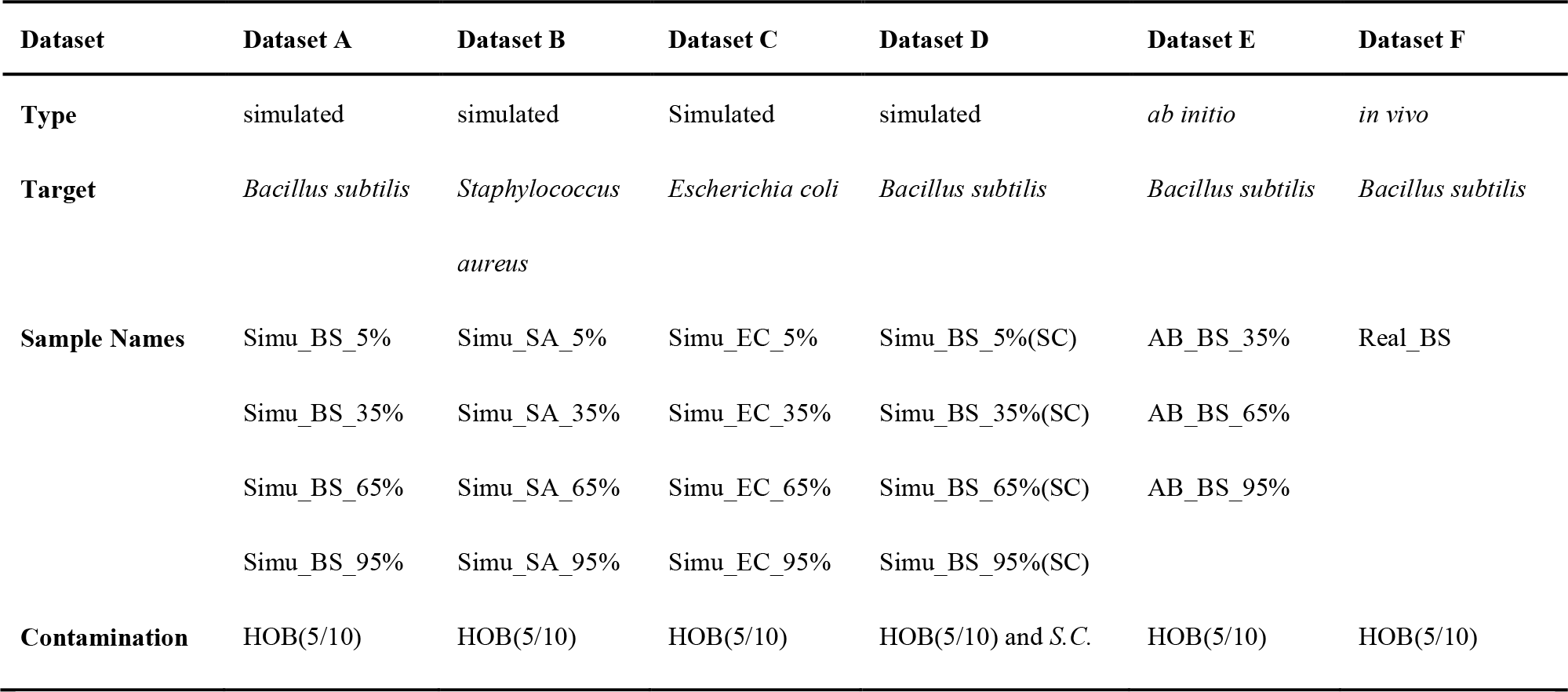
Information about simulated and real metagenomic datasets. Dataset A-D were simulated datasets, Dataset E was *ab initio* dataset, and Dataset F was *in vivo* dataset. Naming style: for simulated datasets, the target species and the relative proportion of reads from target species were provided. For example, “Simu_BS_5%” means that *Bacillus subtilis* was target species, and reads from this target species compose of 5% of all reads in this sample. For *ab initio* datasets, the sample names were defined similarly. The reference genomes of all species were downloaded from NCBI Microbial Genomes website.

| Dataset | Dataset A | Dataset B | Dataset C | Dataset D | Dataset E | Dataset F |
| --- | --- | --- | --- | --- | --- | --- |
| **Type** | simulated | simulated | Simulated | simulated | *ab initio* | *in vivo* |
| **Target** | *Bacillus subtilis* | *Staphylococcus aureus* | *Escherichia coli* | *Bacillus subtilis* | *Bacillus subtilis* | *Bacillus subtilis* |
| **Sample Names** | Simu_BS_5% | Simu_SA_5% | Simu_EC_5% | Simu_BS_5%(SC) | AB_BS_35% | Real_BS |
|  | Simu_BS_35% | Simu_SA_35% | Simu_EC_35% | Simu_BS_35%(SC) | AB_BS_65% |  |
|  | Simu_BS_65% | Simu_SA_65% | Simu_EC_65% | Simu_BS_65%(SC) | AB_BS_95% |  |
|  | Simu_BS_95% | Simu_SA_95% | Simu_EC_95% | Simu_BS_95%(SC) |  |  |
| **Contamination** | HOB(5/10) | HOB(5/10) | HOB(5/10) | HOB(5/10) and *S.C.* | HOB(5/10) | HOB(5/10) |

**Figure 1.**
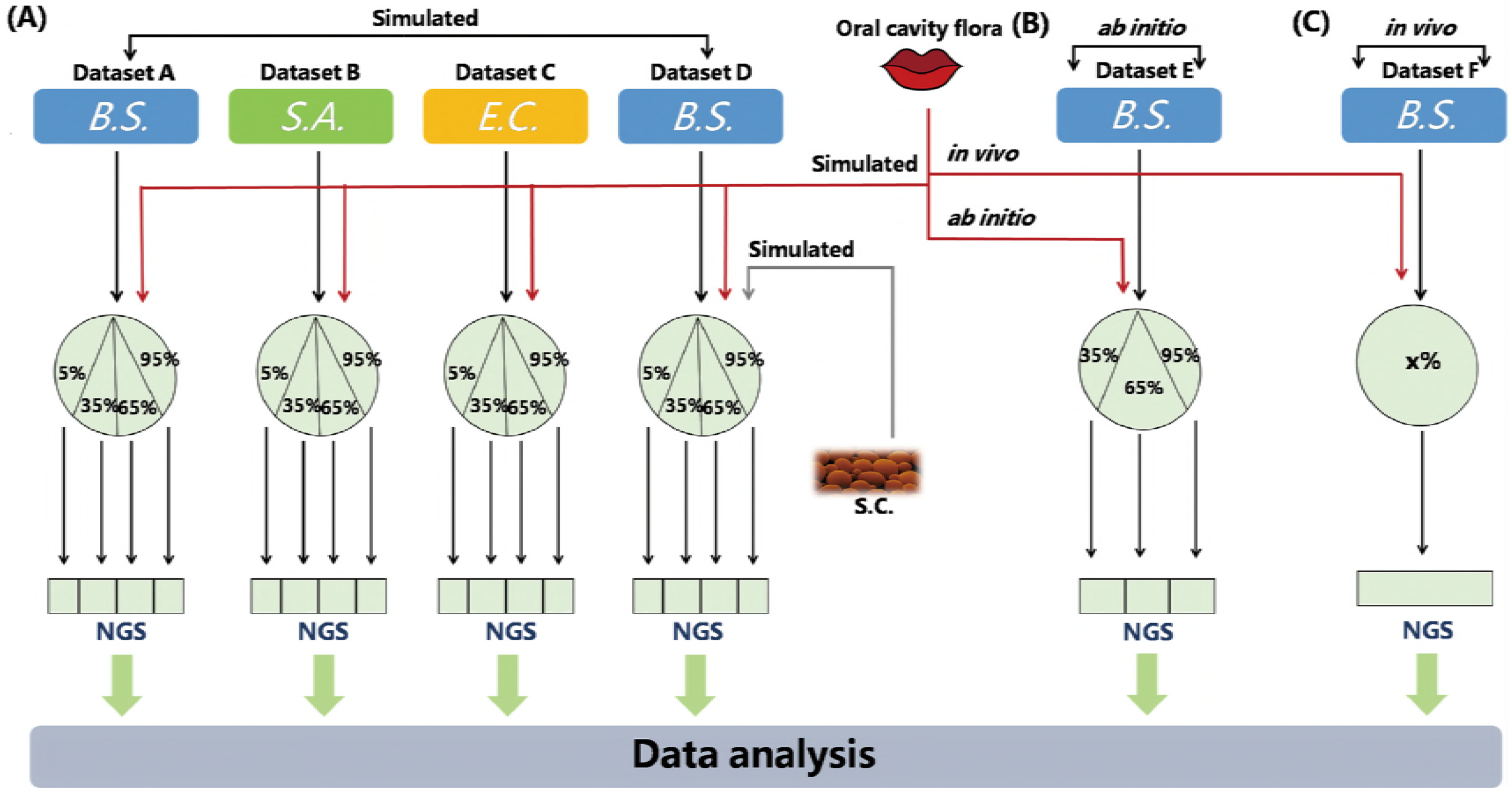
Simulated and real datasets. QC-Blind were tested on three types of data, simulated, *ab initio* and *in vivo* in this study. *B.S., S.A., E.C* represent the three target species (*Bacillus subtilis, Staphylococcus aureus* and *Escherichia coli*.) *S.C.* represents *Saccharomyces cerevisiae*, a source of contamination. The red lines symbolize sequence data from oral cavity flora, while the black lines represent sequence data from target species. (A) In simulated datasets, the target species were mixed with human saliva (or S.C.) at 5%, 35%, 65% and 95%. The grey line represents *Saccharomyces cerevisiae* sequence data. (B) (C)For *ab initio* and *in vivo* datasets, *Bacillus subtilis* (extracted DNA solution and bacteria culture) were mixed with human saliva, the DNA proportions of which were set to 35%, 65%, 95% or unknown ratio (x%).the DNA proportions of which were set to 35%, 65%, 95% or unknown (x%).

#### Metagenomic data simulation

For *in silico* simulated datasets, reads of target and contamination species were generated by NeSSM[15]. In this study, we assume only one target bacteria species in each sample. The target bacteria species used in this study include several model organisms: *Bacillus subtilis, Staphylococcus aureus, Escherichia coli* (dataset A, dataset B, dataset C, **Table 1**). Their reads were mixed with reads generated from the genome of 5 or 10 representative species in human oral microbial community (referred to as HOB(5/10)), which were used as possible human contaminations[16]. Gradient proportions of reads from target species were set to 5%, 35%, 65%, 95%. We also combined *Saccharomyces cerevisiae* with *Bacillus subtilis* and 10 oral bacteria to simulate a special condition with eukaryotic contamination (dataset D, **Table 1**). In each dataset, over 10 million pair-end reads with 100X coverage were generated at the length of 120 bp every 200 bp bin. All other parameters were set as default [15].

#### Ab initio dataset preparation

For *ab initio* datasets, we mixed the real sequencing data of *Bacillus subtilis* with real metagenomic sequences from human saliva samples (dataset E, **Table 1**), with the relative proportion of reads from target species (*Bacillus subtilis*) set at 35%, 65%, 95% for different datasets. These samples were named AB_BS_35%, AB_BS_65%, AB_BS_95% respectively. We did not perform any filtration of human contaminations from saliva samples (nearly 1,000 species in human saliva in 2014) [16], in order to maintain them in their natural state, even though the lack of filtration may adversely affect the contig assembly and clustering process.

For both *ab initio* and *in vivo* dataset preparation, real DNA extraction and sequencing are needed. Their procedures are detailed in sub-section “*In vivo* sample preparation, DNA extraction and sequencing”.

#### In vivo sample preparation, DNA extraction and sequencing

The *in vivo* datasets used in this study were metagenomic (not 16s rRNA) datasets from real community samples as prepared below: after being activated, *Bacillus subtilis* 168 was cultured overnight till its OD_600_ value reaches between 0.6 and 0.8. All the *Bacillus subtilis* were centrifuged at 12000 rev min −1 (12114g) for the following experiment. The fresh saliva was collected from three healthy adults without drinking water or gargling for about 30min before sample collection. Then 200ul fresh saliva was added to the *Bacillus subtilis* culture before DNA extraction, amplification and sequencing. This sample was named Real_BS (dataset F, **Table 1**).

Modified CTAB method[17–19] was chosen for obtaining high MW metagenomic DNA of samples. 5 ml lysis buffer (Cetyl Trimethyl Ammonium Bromide, 1% w/v; EDTA, 100mM; NaCl, 1.5mol l ^−1^; Sodium phosphate, 100mmol l ^−1^; Tris-Cl pH 8.0, 100mmol l ^−1^) and 20 μl Proteinase K was added into 15 ml liquid sample (*B.S.* culture or its mixture with saliva), followed by gentle shaking at 100rev min ^−1^. SDS was added to a final concentration of 1% and the reaction was incubated at 65°C for 30min with intermittent shaking. After the above steps, an equal volume of saturated phenol, chloroform and isoamyl alcohol (25: 24:1) was added to the mixture and centrifuge at 12000 rev min −1 (12114g) for 10min to collect supernatant that are free from protein. Then, this step is repeated once. Metagenomic DNA was precipitated with 0.6 volumes of isopropanol for 30min at −20 °C and pelleted by centrifugation at 12,000rev min −1 (12114g) for 10min. DNA was washed twice with 70% ethanol and finally dissolved into a 200μl of TE (1X), pH 8.0. The genomic DNA of *Bacillus subtilis* 168 and the mixture of *Bacillus subtilis* 168 with human saliva were extracted with Soil Genomic DNA kit(CWBIO).

Before sequencing with Illumina Miseq-2000, DNA samples were quantified using a Qubit^®^ 2.0 Fluorometer (Invitrogen, Carlsbad, CA) and its quality was checked on a 0.8% agarose gel. 5-50 ng metagenomic DNA in high quality was used as the template for amplifying the V3-V4 diameter region of 16S rRNA genes for each individual sample, with “5′-CCTACGGRRBGCASCAGKVRVGAAT-3′” used as the forward primer and “5-GGACTACNVGGGTWTCTAATCC-3′” as the reverse primer. The sequencing library was constructed using a MetaVxTM Library Preparation kit (GENEWIZ, Inc., South Plainfield, NJ.USA). Then indexed adapters were added to the ends of 16S rDNA amplicons by limited-cycle PCR. Verified by Agilent 2100 Bioanalyzier(Agilent Technologies, Palo Alto, CA, USA), and quantified by Qubit^®^ 2.0 Fluorometer (Invitrogen, Carlsbad, CA) and real-time PCR(Applied Biosystems, Carlsbad, CA, USA), the DNA libraries were normalized for sequencing. All sequencing reactions were performed on the Illumina MiSeq platform using paired-end sequencing technology (2*300 bp).

### 2. Analytical procedure

An overview of our quality control pipeline is shown in **Figure 2(A)**. First, real sequencing data are trimmed by Trimmomatic-0.36 to remove low quality bases and reads[20]. 3 leading/trailing bases are cut if their quality scores are below a quality threshold. Reads with lengths that are too short (μ50bp as default) are also discarded. 16s rRNA genes are extracted from remaining reads, for species identification and quantification. Then read assembly, contig binning and marker gene mapping are performed in order.

**Figure 2.**
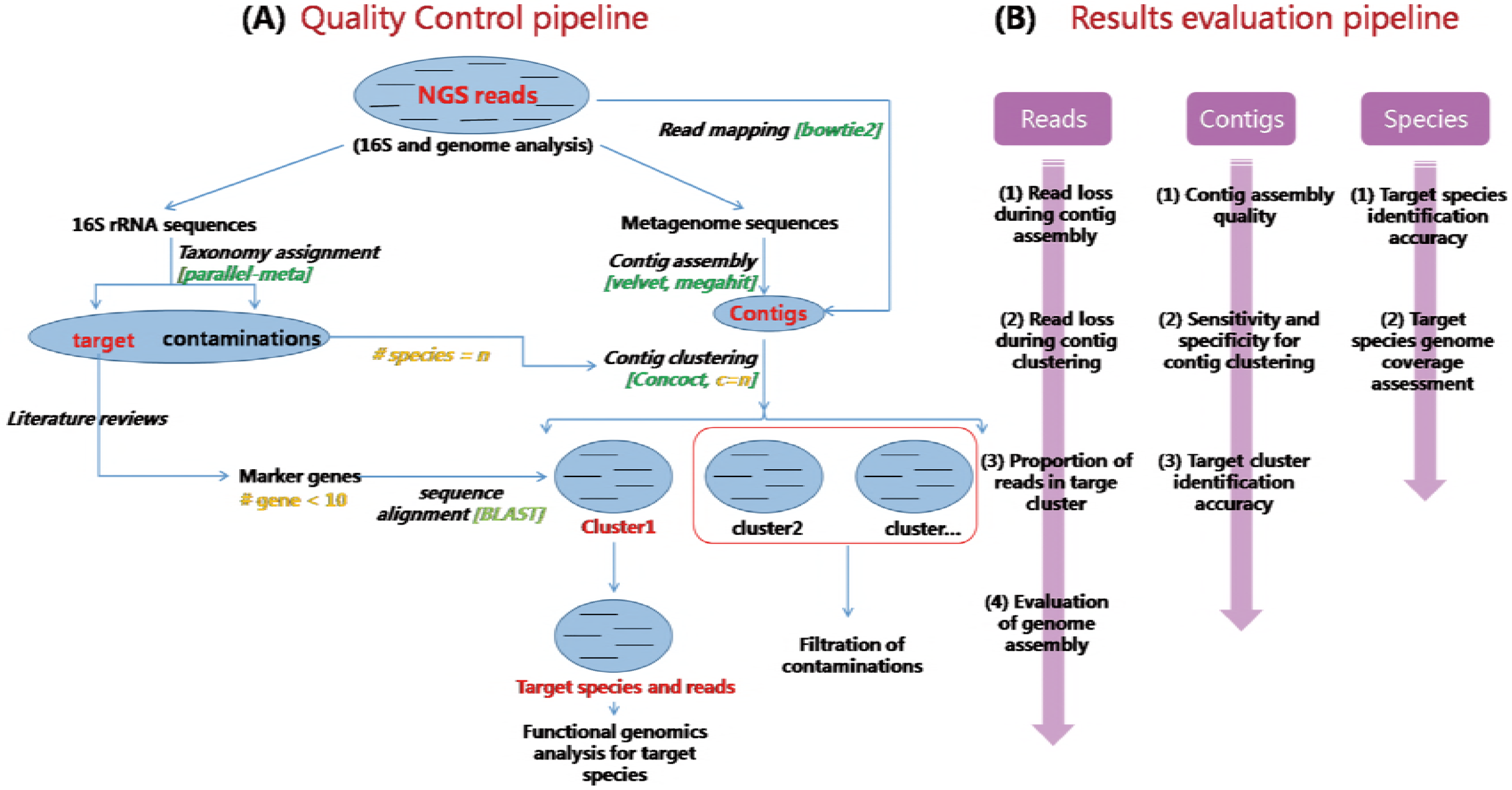
Schematic demonstration and evaluations of QC-Blind. (A) An overview of the QC-blind pipeline. Based on NGS data, on one hand, 16s rRNA genes were extracted and compared with bacteria 16s rRNA database to count species number; on the other hand, raw reads were assembled into contigs, binned into species-level groups, and then target clusters were identified through mapping marker genes of target species onto them. Among these steps, the marker gene mapping step could finally determine which cluster of contigs and reads belong to the target species. (B) Evaluations for QC-blind were performed at three levels, read, contig and species. For clustering quality, the purity and concentration of target and other clusters were measured. For contamination removal, sensitivity and specificity were calculated. For data loss, target reads and contigs fail to pass at each step were counted. For functional analysis, the coverage of target genome at base and gene level were calculated.

#### Identification of target and contamination species

The taxonomical profiles were generated by the Parallel-Meta pipeline (version 2.0)[15]. 16s rRNA sequences were extracted from raw sequencing data through an HMM. These sequences were searched against the Greengene database to identify their species.

The total number of species identified was used as input at a contig binning step, which aims to provide better accuracy for clustering. An additional eukaryotic18S rRNA database was used as reference when processing the dataset with *Saccharomyces cerevisiae*. Choosing the number of clusters become difficult for unknown contaminants whose information are not recorded in 16s RNA or 18S rRNA database, but this approach is practical and performs well for target identification and contamination filtration.

#### Assembly of contigs from community data

Two assemblers were applied to assemble contigs from community reads. One of the assemblers selected was Velvet[15], which could deal with *de novo* genomic assembly and short sequencing reads alignment. For Velvet, we use the *velveth* command to construct the data set as preparation work, and *velvet* command to build the *de Bruijn* graph from the *k-mers* obtained by *velveth* and extract the contigs. We use 12 for *k*, and set other parameters to auto or default. The other one was MEGAHIT[21], an assembler designed specifically for complex metagenomics via succinct *de Bruijn* graph. It is worth mentioning that by using these two assemblers, the abundance information has been intrinsically taken into consideration.

For simulated metagenomic datasets, assembly was performed on two assemblers to compare their performance. Basic assembly statistics were extracted and compared. As MEGAHIT has been shown to be superior to Velvet through the analysis of simulated data, only MEGAHIT was used to process *ab initio* and *in vivo* dataset.

#### Contig binning with CONCOCT

Contig binning is the central step of QC-Blind. Out of all the existing and evaluated binning algorithms, CONCOCT is selected because: first, both sequence composition and coverage across multiple samples were considered in contig binning; second, it could handle both single sample and multiple samples. These make CONCOCT a suitable choice for batch process of possibly contaminated samples [10]. For multiple species, co-assembly is a necessary prior to running CONCOCT, as CONCOCT takes contigs as input to maximize the number of genomes that could potentially be resolved. Contigs are limited to lengths from 1,000 bp to 10,000 bp, the lower limit filters low quality contigs while the upper limit cuts fragments for more statistical weight. The number of clusters was precisely determined with 16s rRNA method by Parallel-Meta[22]. Contigs would be clustered into species-level groups after the processing of CONCOCT. Again, we have to emphasize that by using CONCOCT, the abundances (read depth) of contigs in each of the clusters would become similar.

#### Marker gene selection and mapping

Utilizing marker gene for target species cluster identification helps us overcome the situation where we do not have the complete or partial reference genome, except for a few marker genes. This is realistic for most of the contemporary sequencing tasks, in which we only know a few of the targets’ marker genes [23, 24].

Marker genes were generated by the combination of MetaPhlAn2[14] and manual curation. The clusters generated in the previous step were mapped to marker genes of target species by BLAST (e-value cutoff = 1e-20). Except for a few well-studied species that have had their whole genome sequenced, we have knowledge on only a few genes for most of the other species, such as important regulators in their metabolism, proliferation, or 16s rRNA genes. We use these genes as markers to identify contig clusters that belong to the target species. The number of usable marker genes depends on the knowledge we have. The more unique the genes, the more specific the identification would be. Through MetaPhlAn2 selection and consulting literature, q-PCR markers ftsZ, lytF, nsrR, spo0A, ygxB, yjbH, yjbI were selected for *Bacillus subtilis*[8, 25, 26], acpP, casA, cof, dxs, fabB, fabF, leuO, tesA, uidA were chosen for *Escherichia coli*[27–31].

Contigs containing marker genes for target species are identified as belonging to the target species (defined as target contigs). Based on these assignments, raw reads were mapped to contigs identified as belonging to the targets with BOWTIE2[32] (defined as target reads). Statistics of total reads number and target reads number in every step could then be evaluated. For *ab initio* and *in vivo* datasets, only the target reads or contigs were classified through mapping to reference of *Bacillus subtilis*, since it was impractical to classify each read of contaminations to their source species, especially when many of them do not yet have their whole genome sequenced [16].

### 3. Evaluation methods

Our assessment of QC-Blind is based on the purity of the clusters, target distribution, sensitivity, specificity, data loss and coverage (**Figure 2(B)**).

#### Purity of clusters

DS (dominant species) is defined as the species whose contigs (and reads) outnumber other species’ in the cluster. In each cluster, all the contigs were mapped to their reference genome database consisted of both target bacteria and contaminations by BLAST, to identify their source species. Reads were mapped to contigs by BOWTIE2[32] and thus inheriting taxonomical information from contigs. Purity was defined as the proportion of contigs or reads of the DS in each cluster, after contig binning through CONCOCT[10] (**Formula 1** and **Formula 2**).

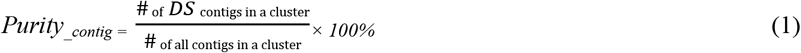

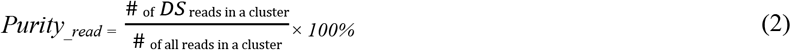

Purity of each cluster was categorized and evaluated at three levels, 100%, 90%+, 80%+. The proportion of clusters exceeding these thresholds among all the clusters reflects the quality of the clustering.

#### Target contig and target read concentration and distribution

We define clusters that contain contigs (and reads) from target species (TS) as target clusters (TC), no matter how many contigs are in TC. The proportion of target contig or target read in each clusters, as well as their distributions among all target cluster are measured in **Formula 3** and **Formula 4**.

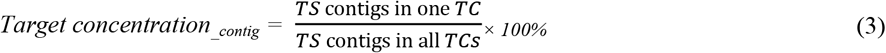

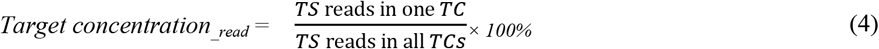

Target concentrations in all clusters would thus form a distribution, which is named “Target distribution”. A more biased target distribution indicates a better clustering result, and vice versa.

#### Sensitivity and specificity

“Target/Contamination Dichotomization” were also measured by sensitivity and specificity, which provided quantitative information about contaminations correctly removed and target sequences successfully preserved (**Formula 5-8**). Here, the contigs or reads of target species in the clusters found by marker genes alignment were considered as true positive (TP), while those of other species were considered as false positive (FP). The contigs or reads mapping on the target genome were considered as ground truth (GT), which contained those identified or unidentified contigs or reads in target clusters.

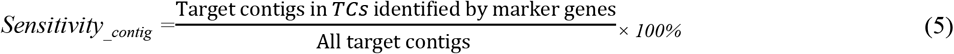

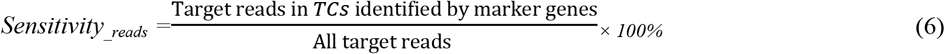

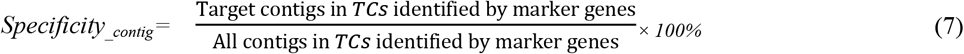

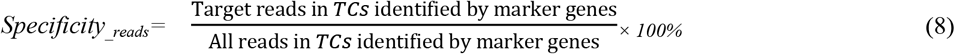

#### Coverage evaluation

We performed functional evaluations to examine how much of the target bacteria species sequencing data has been kept after the quality control process, and whether they retain the functional genomics of the target species. The coverage in target genome was calculated at both base and gene level (**Formula 9-10**). At the base level, the number of bases in the genome that have been mapped for one or more times was measured (mapped base, MB) and compared with the length of the whole genome (total base, TB). Statistical data were summarized after target reads alignment. At the gene level, genes covered by contigs in our target clusters were viewed as mapped genes (MG), while the genes for the all species were considered as total gene (TG). Genes preserved after the processing were identified by the *intersect* command of BEDTools[33]. Obviously, gene and base coverage for unprocessed simulated data were both 100%.

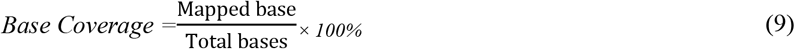

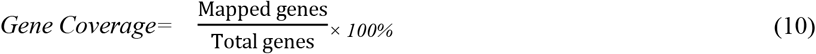

## RESULTS AND DISCUSSIONS

Simulated metagenomic datasets, each consisting of 4 samples with different multi-species complexity, were selected to benchmark the performance of QC-Blind for target species identification (**Figure 1**). Here we present the results for dataset consisting of *Bacillus subtilis* as target species (5%, 35%, 65% and 95% reads proportion) and 10 human oral bacteria as contaminations [16] (**Table 1**). Results for target species of *S.A.*, *E.C.*, and *B.C* mixed with *S.C* were shown in **Supplementary File 1**.

### Target and contamination species identification

Target species can be completely identified at genus level, along with 88% of contaminations identified on average (**Supplementary File 2-Table 1**). The number of species acquired in this step could increase the purity of clustering, although the unsupervised contig binning method utilized in QC-Blind does not necessarily require this parameter[10]. However, it was also found that most eukaryotic organisms (usually not contaminations) would not be identified at species level by this method.

### Read assembly and contig binning

For read assembly, results based on Velvet[34] and MEGAHIT[21] were compared, showing that MEGAHIT would generate less contigs with longer N50 (eg. For BS 5%, 603 contigs were generated with N50=154200 by MEGAHIT, while 10667 contigs were assembled by Velvet with N50=39369), thus we expect contigs from MEGAHIT to outperform those from Velvet in downstream analysis (**Supplementary File 2** | **Table 2**).

Purity of cluster evaluation indicated that most of these contigs generated by QC-Blind were dominated by single species. At the read level, 59.6% (28 of 47) of the clusters reached 90% purity on average, and 72.3% clusters reached 80% purity (34 of 47) (**Figure 3(A)**). At contig level, more than 50% of the clusters reached 90% purity. For target concentration, each dataset had a single main cluster that contained over 94% target contigs. Noticeably, the dominant clusters in three simulated datasets were of 100% purity, except that purity of Simu_BS_95.0% dominant cluster 8 was 95% (**Figure 3(B)**).

**Figure 3.**
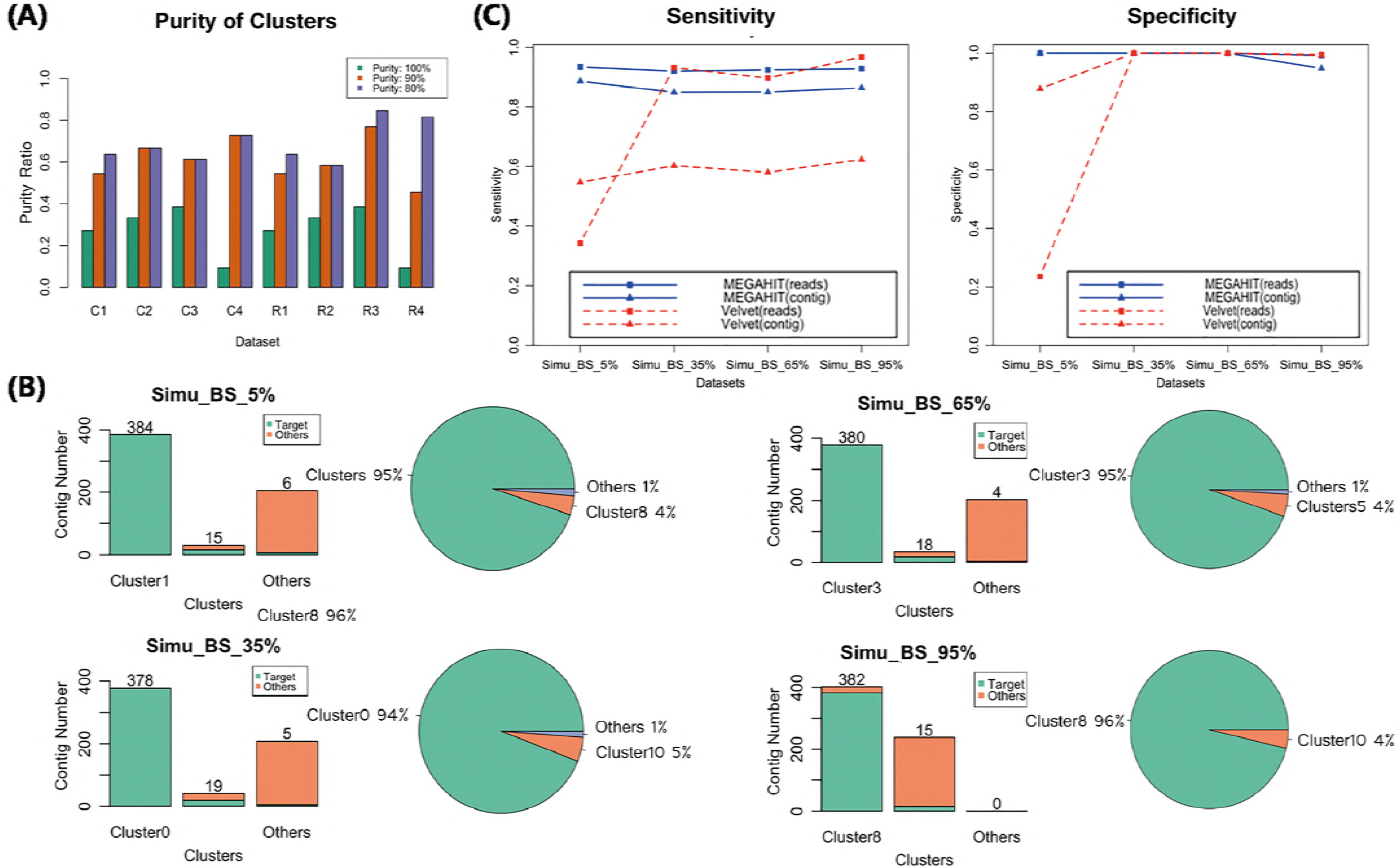
Evaluation of contig binning and contaminations removal. (A) 11, 12, 13, 11 clusters were generated respectively for the four datasets with target reads fraction of 5%, 35%, 65%, 95%. Bar plots shows the purity ratio of reads and contigs in all clusters in each sample. C1 – C4 and R1– R4 stands for contigs and reads of the four samples, respectively. Purity were measured at 100%, 90%+, 80%+ level, in which green lump represents 100% pure, the orange lump represents purity over 90%, the blue lump represents purity over 80%. (B) Target distribution and concentration in four samples. Clusters with the largest and second largest number of target contigs were shown, combined with all other clusters that has at least one target contig. The green lump represents target species, while the orange lump represents other species. Distribution of all target contigs were also shown in pies.(C) Sensitivity and specificity of the results of the four groups at MEGAHIT-BLAST (blue lines) and Velvet-BLAST (red lines) pipeline. The squares stand for results based on read level and the triangles stand for results based on contig level.

Taken together, this contig binning method could resolve single highly concentrated and pure target cluster from multiple species. Considering possible artifacts produced during read mapping on the simulated datasets, we anticipated that the method would actually perform better for real datasets.

### Evaluation of sensitivity and specificity for target species read assignment

In general, the sensitivity and specificity values for target species read assignment of MEGAHIT-processed data were both high (**Figure 3(C)**). Sensitivity values were on average 92.7% in four samples, while specificity values of those were even higher for both target contigs and reads: 100% assignment specificity in Simu_BS_5%, Simu_BS_35% and Simu_BS_65%, showing that the target information in target cluster can be extracted with very few contaminations remaining. However, the sensitivity and specificity evaluation of Velvet-processed data were extremely low at the dataset with 5% target reads (34.3%, compared to 93.5% in MEGAHIT), which raised question on the ability of Velvet to deal with severely contaminated data. Velvet’s sensitivity at the contig level was also not optimistic.

Taken together, the evaluation of sensitivity and specificity for target species read assignments showed the superiority of using MEGAHIT in QC-Blind method. Thus, in the following analyses, we adopted MEGAHIT in the QC-Blind method as default.

### Data loss in screening process

The information loss that we generally experience as the target information progress from raw reads, to read assembly, contig binning, and then marker gene mapping, decreases as the proportion of target species increases (**Figure 4(A)**). The greatest data loss on read level occurred in marker gene mapping. the proportion of reads loss were up to 5.31%, 5.91%, 5.32%, 4.87% for each simulated dataset with target fraction from 5% to 95%. Samples without a dominant species (e.g. Simu_BS_5%, Simu_BS_35%) encountered difficulty on assigning all reads correctly, as there may not be enough unique reads from them to reconstruct a complete genome for species identification[10]. Some short contigs were also filtered out during contig binning with proportion of 6.47%, 9.66%, 10.07%, 9.05% respectively (**Figure 4(B)**). Data loss at read level is almost negligible (less than 2%) in reads assembly and contig binning (**Figure 4(C)**), implying that the QC-Blind method has the capacity to preserve genetic information hidden in unique reads during marker gene mapping.

**Figure 4.**
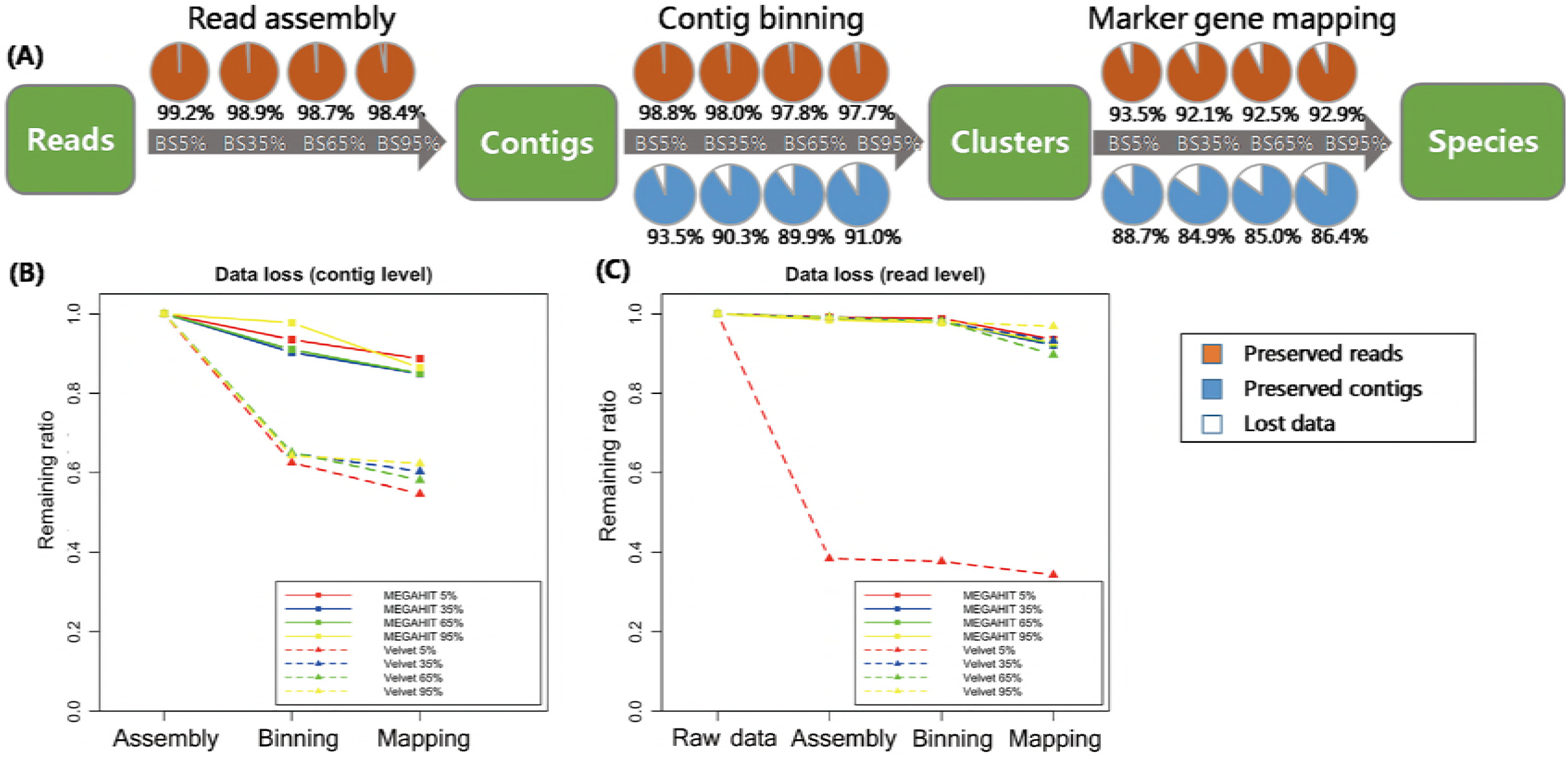
Data loss assessment for simulated data. (A) Percentage of target contigs and reads preserved in each step (read assembly, contig binning, marker gene mapping), for MEGAHIT pipeline. The four circles in a row represent groups in which the frequency of target species increases from 5% to 95%. The orange lumps represent preserved reads, the blue lumps represent preserved contigs, while the white lumps represent lost data. (B) Comparison of Data loss in the steps of read assembly, contig binning and marker gene mapping between MEGAHIT-BLAST (full squares) and Velvet-BLAST (dot lines and triangles) pipeline at contig level. The red, blue, green and yellow lines represent the samples with proportion of target species from 5% to 95%. (C) Data loss comparison between MEGAHIT-BLAST (full squares) and Velvet-BLAST (dot lines and triangles) pipeline at read level. The red, blue, green and yellow lines represent the samples with proportion of target species from 5% to 95%.

By contrast, analysis results on the data loss issue by Velvet was unsatisfactory, indicating its inability to deal with highly contaminated data. At contig level, over 35% contigs were lost after contig binning and mapping of Velvet-processed data in all four samples. At read level, Velvet performed comparably as MEGAHIT in three samples, except that in Simu_BS_5%, 61.6% reads were lost after contig assembly by Velvet.

### Base and gene coverage analysis

QC-Blind results have nearly perfect coverage of genomic information regardless of the different complexities of the samples. Through mapping these reads back to the genome, on average 93.5% of the bases could be covered for one or more times, indicating the potential of this method to reconstruct a complete genome (**Figure 5(A)**). More notably, this coverage is consistent among the four samples (94.1%, 92.9%, 93.2%, 93.8%), which suggests that QC-Blind is able to work with samples of different complexities. Similarly, most of the annotated genes are also covered with processed reads (on average 93.8%). The Arginine biosynthesis pathway, a critical pathway for *Bacillus subtilis*’s metabolism[35], was selected as an example for examination(**Figure 5(B)**); all the 20 genes that are involved in this pathway could be found in the processed reads (after quality control) of the four datasets (**Figure 5(C)**).

**Figure 5.**
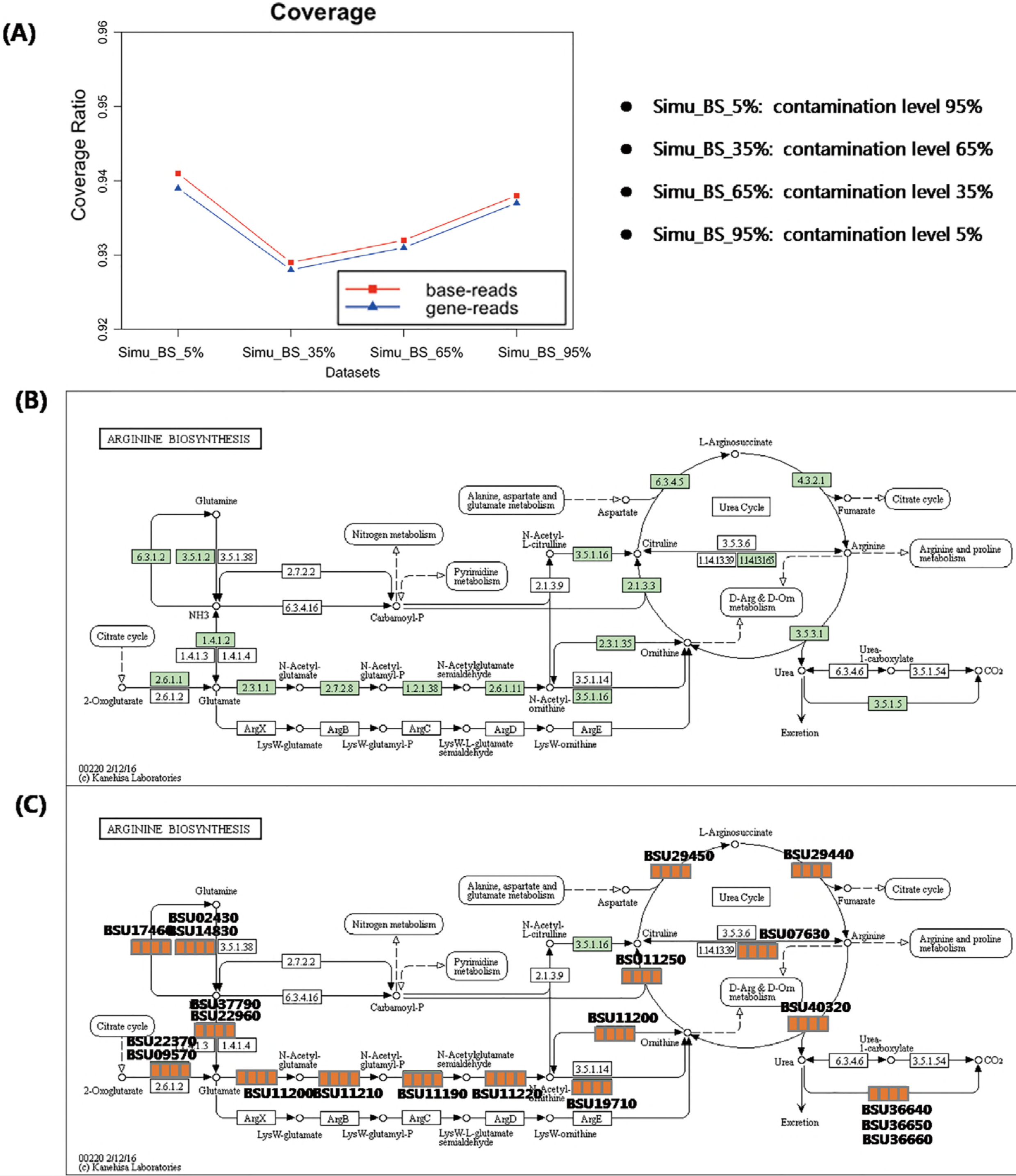
Coverage and pathway reconstruction based on simulated data. (A) Y-axis represents base coverage (red line, squares) and gene coverage (blue line, triangles) of target genome as described in Methods. X-axis stands for four samples with target species from 5% to 95%. (B) Arginine biosynthesis pathway of *Bacillus subtilis* from KEGG. (C) Genes successfully found in processed data marked with orange squares, and four squares stands for each sample, in which yellow and white stands for exist and non-exist.

The above assessments of QC-Blind based on simulated data have not only demonstrated the possibility but also the high fidelity of the reference-free QC. This performance can be attributed to the fact that the vast majority of target contigs were binned into a single cluster, which makes it more convenient for the identifications of marker genes. Certainly, the selection of marker genes was very crucial, as their uniqueness among microbial community would assure high specificity in target/contamination classification.

Hence for simulated data, the resultant near-perfect coverage proved that the additional work performed on screening is worthwhile. We have put other simulation settings and results in **Supplementary File 1** including analytical results of Dataset B~D (**Figure 1(A)**).

### Analysis based on ab initio and in vivo datasets

Before bacterial contamination screening, QC-Blind was able to capture the genetic information of target species in *ab initio* and *in vivo* datasets that were contaminated by large proportion of human-oriented reads (416, 679, 339 reads were identified from bacteria floras of human saliva in this study). This might affect the contig assembly and clustering process.

Single dominant cluster for the target species *Bacillus subtilis* were successfully determined in each *ab initio* and *in vivo* dataset, with cluster 34 in AB_BS 35% containing 99.6% target reads, cluster 33 in AB_BS 65% containing 99.9% target reads, cluster 32 in AB_BS 95% containing 99.5% target reads and cluster 14, 65, 78 together containing 59.7% of target reads (**Figure 6(A)**), while a lot of contigs from contamination species with very few sequences were not classified into independent cluster in AB_BS 65% and AB_BS 95%. All three dominant clusters were identified by marker genes with high specificity (**Figure 6(B)**). However, the sensitivity of AB_BS 65% at read level dropped to 47.5% while the sensitivity of AB_BS 35% and AB_BS 95% and Real_BS remained high.

**Figure 6.**
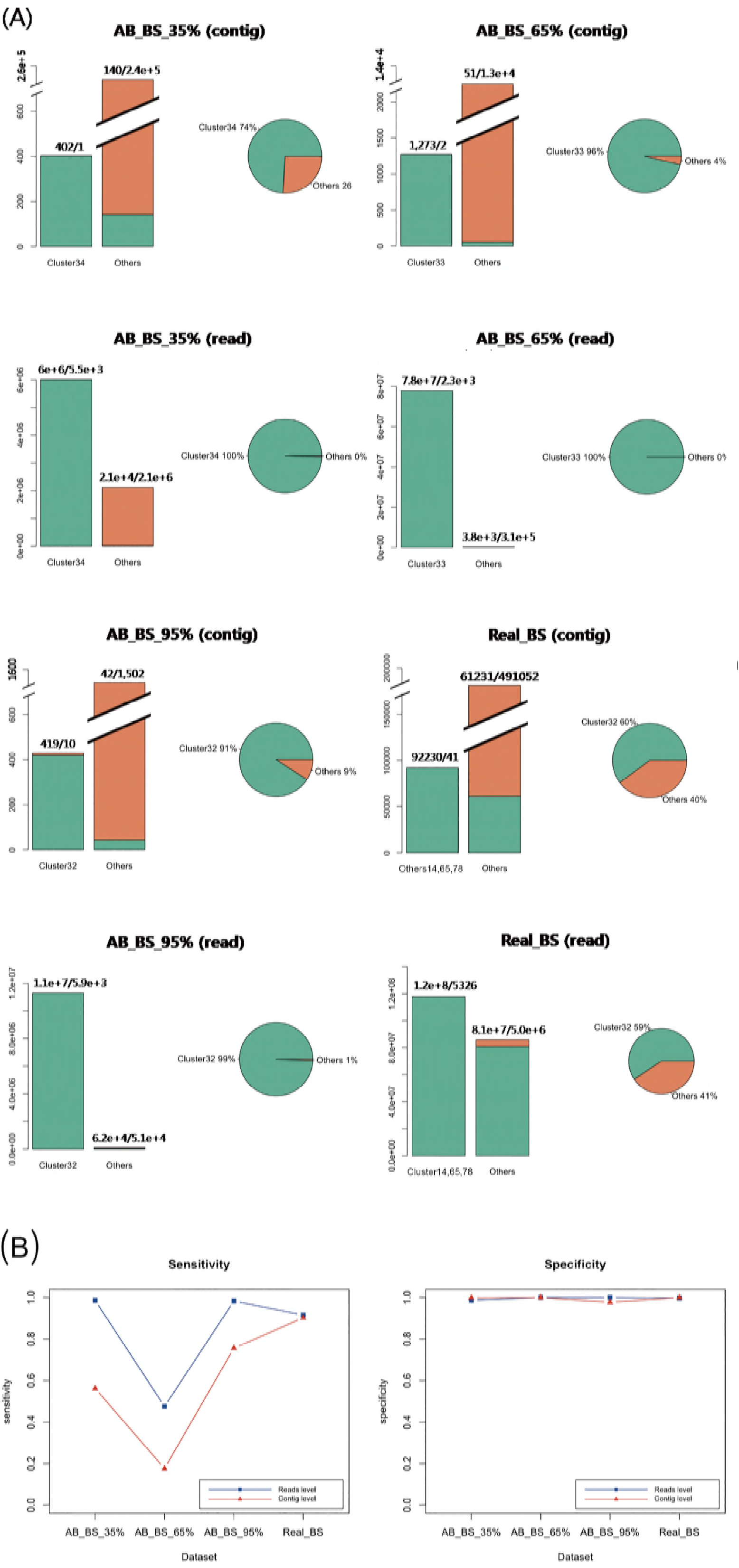
Evaluation of contig binning and contaminations removal for real data. (A) Target distribution and concentration in four samples. In Barplots, clusters with the largest number of target contigs were shown, combined with all other clusters that has at least one target contig. The green lump represents target species, while the orange lump represents other species. Distributions of all target contigs were also shown in pies. The green lump stands for the cluster with the largest number of target contigs, while the orange lump stands for other clusters containing target contigs. The clusters are marked by original cluster ID from QC-blind pipeline (e.g. Cluster34). (B) Sensitivity and specificity of the results of the three groups at MEGAHIT-BLAST. The blue lines with squares stand for read level results and the red lines with triangles stand for contig level results.

For data loss ratios, our method’s performance on read datasets remained high level (**Figure 7(A)**), except that less than 20% contigs remained in AB_BS 65% with less than 30% reads after assembly (**Figure 7(B)**). For possible explanation for this abnormal phenomenon, we found that the N50 and average contig length of AB_BS 65% were the lowest among the three (**Table 2**), a large number of its contigs were filtered due to the 600bp cutoff of CONCOCT. This stringent cutoff was set to remove low quality reads and keep specificity at high level, which if set lower, might recover the data loss (**Supplementary File 3**). No more than 6% of reads were lost in marker gene mapping.

**Figure 7.**
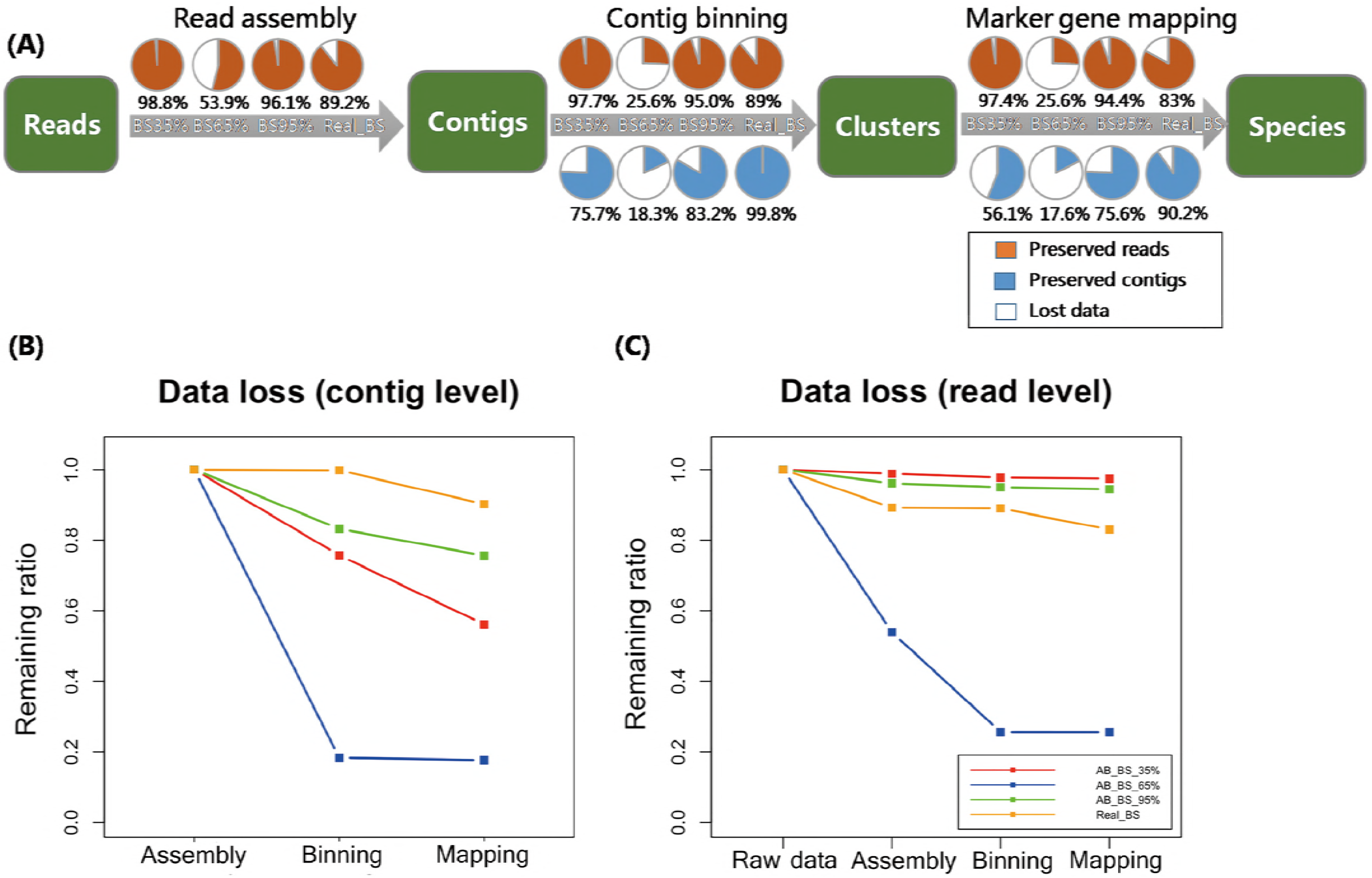
Data loss assessment for real data. A) Proportion of target contigs and reads preserved in each step (read assembly, contig binning, marker gene mapping), for MEGAHIT pipeline. The four circles in a row represent groups in which the frequency of target species increases from 35%, 65% to 95%, as well as real BS sequencing data. The orange lumps represent preserved reads, the blue lumps represent preserved contigs, while the white lumps represent lost data. (B) Data loss in the steps of read assembly, contig binning and marker gene mapping of MEGAHIT-BLAST pipeline at contig level. The red, blue, green and yellow lines represent the samples in which the frequency of target species increases from 35%, 65% to 95%, as well as real BS sequencing data.(C) Data loss of MEGAHIT-BLAST pipeline at read level. All results were calculated based on simulated dataset A.

For base and gene coverage, the performance of QC-Blind on real datasets was consistent with that on simulated datasets, as the analytical results of AB_BS 35%, AB_BS 95% and Real_BS datasets were all over 98%, except for AB_ BS 65%(**Figure 8(A)**). The results reaffirmed the potential of QC-Blind to reconstruct genome from real sequencing data with contaminations (**Figure 8(B), (C)**).

**Figure 8.**
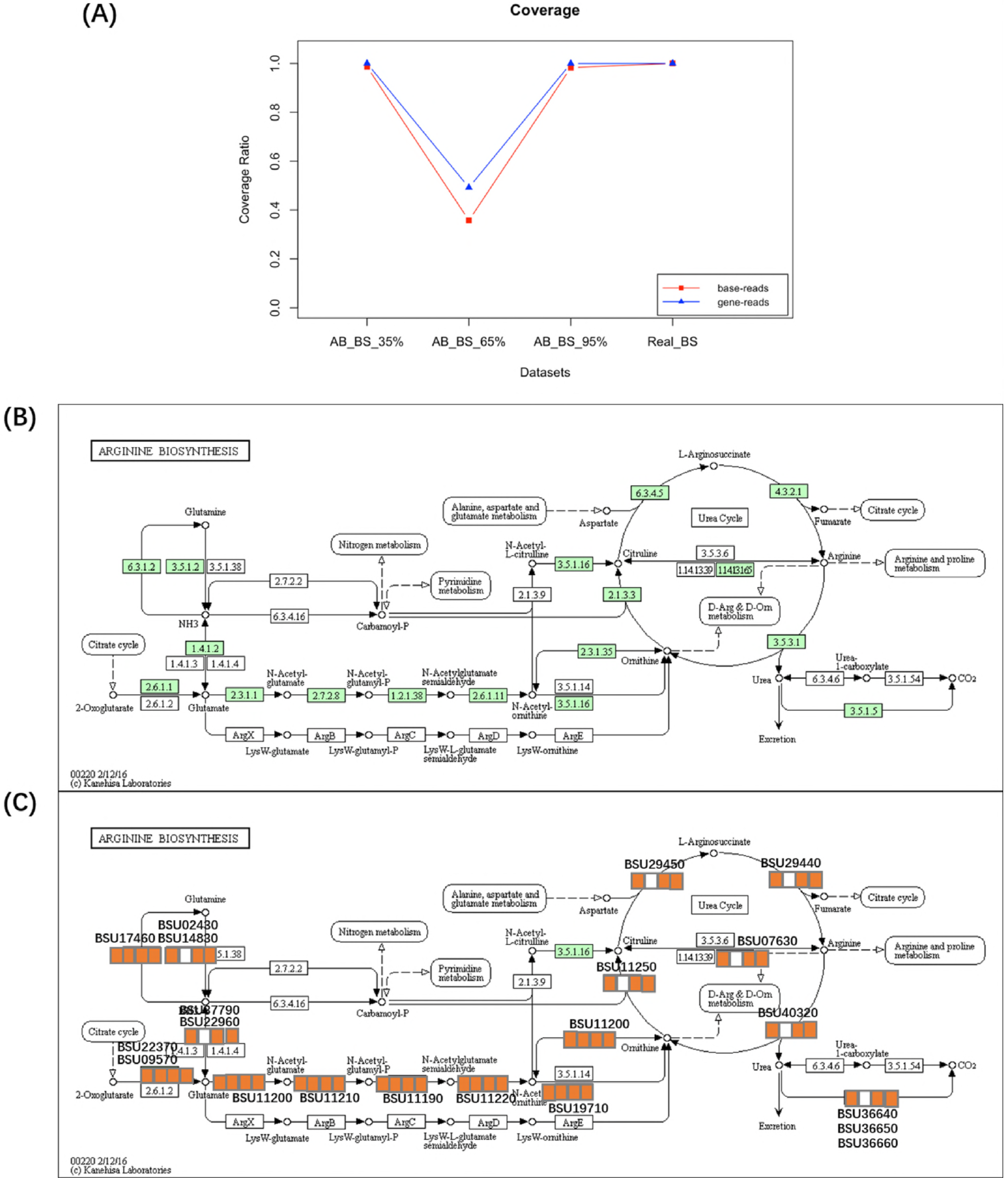
Coverage and pathway reconstruction for real data. (A) Y-axis represents base coverage (red line, squares) and gene coverage (blue line, triangles) of target genome as described in Methods. X-axis stands for four samples with proportion of target species from 5% to 95%. (B) Arginine biosynthesis pathway of *Bacillus subtilis* from KEGG. (C) Genes successfully found in processed data marked with orange squares, and three squares stands for each sample, in which yellow and white stands for exist and non-exist.

**Table 2.**
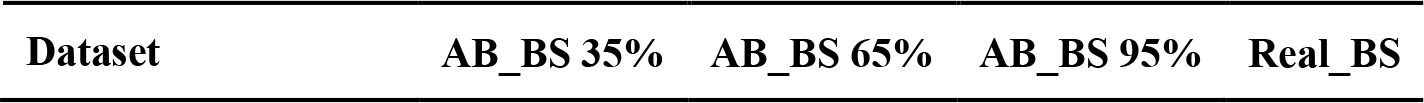
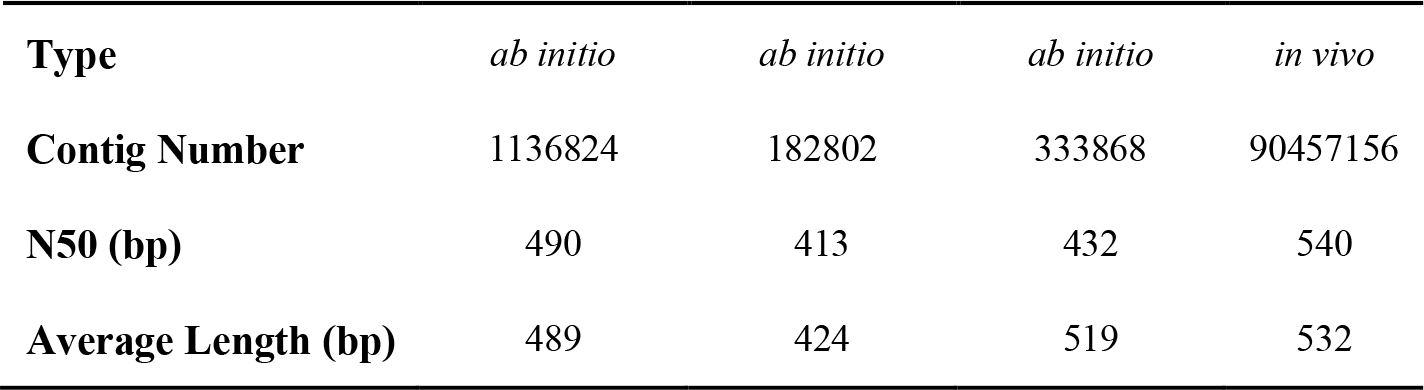
Assembly result summary for *initio* and *in vivo* datasets. Contig number, N50 and average length of contigs assembled by MEGAHIT from real datasets are shown. More information of AB_BS_35**%,** AB_BS_65**%,** AB_BS_95**%**and Real_BS are shown in **Table 1**.

| **Type** | *ab initio* | *ab initio* | *ab initio* | *in vivo* |
| --- | --- | --- | --- | --- |
| **Contig Number** | 1136824 | 182802 | 333868 | 90457156 |
| **N50 (bp)** | 490 | 413 | 432 | 540 |
| **Average Length (bp)** | 489 | 424 | 519 | 532 |

### Comparison with existing QC tools

There is no existing reference-free QC tool similar to QC-Blind up till now, so it is difficult to conduct a completely fair performance comparison with QC-Blind. However, it is still valuable to compare QC-Blind with commonly used QC tools in terms of contaminant identification and filtration.

Kontaminant is a kmer-based Contamination screening tool [36, 37], which has proved to be effective in host filtering for novel viral discovery(ref3). Compared to QC-Blind, Kontaminant is limited by the completeness of existing k-mer database, which lead to its inability to work on unknown contaminations.

Another general QC analytical tool is FastQC, which is able to generate a comprehensive report on the quality profile of the reads [37, 38]. With respect to contaminant identification, it could detect the overrepresented sequence through assessing per base sequence content, per base GC content and per sequence GC content, which can be used as evidence that the library is contaminated[38]. Although FastQC is able to detect unknown contaminations, it cannot remove contaminants well, especially in situations where there are multiple contaminating species. Compared to FastQC, QC-blind implements species-based contigs binning method (ideally one cluster is from one species), which performs better in the case that the contaminant is a mixture of multiple species.

QC-Chain [12] has good performance in identifying and removing contaminations. However, the false positive rate remains high. Compared to this method, the purity of target clusters in QC-Blind is much higher, and marker genes can map the target clusters accurately, which could ensure higher true positive rate.

### Efficiency evaluation

QC-Blind ran in less than 12 hours on a single processor on datasets with 40 million paired-end reads, with varying time depending on the sequencing quality and contig number. The greatest proportion of time was consumed on contig binning. Due to the time complexity of the clustering algorithm, the running time increased significantly when working with larger number of contigs, which result from the use of lower cutoff contig lengths. Thus, a reasonable cutoff, an improved clustering algorithm, as well as the utilization of multiple processors could be taken into consideration to reach higher efficiency.

## CONCLUSION

In this study, we proposed the QC-Blind pipeline for assigning reads to target species. QC-Blind first uses the well-established 16s rRNA approach for species identification to identify the number of species. It then employs read assembly (Velvet and MEGAHIT) and contig binning (CONCOCT) sequentially to cluster the contigs and reads. Finally, it uses the marker genes to identify the contig clusters.

A systematic assessment of QC-Blind based on *in silico*, *ab initio* and *in vivo* datasets with different fraction of contaminations, showed QC-Blind to perform well on screening microbial contamination. After processed by QC-Blind, contigs and reads were highly concentrated in pure clusters and were easily identified through their marker genes. The clusters are almost homogeneous even when samples are contaminated by more than 50% heterogeneous reads. The reads which QC-Blind recovered consists of a large proportion of the genome of the target species. However, the performance of QC-Blind on simulation datasets, *ab initio* mixture of species and *in vivo* real datasets showed certain differences: data loss was more significant in a few real datasets such as BS 65%.

Our tests show that QC-Blind is able acquire high-quality sequencing data, as well as reduce the widely presented “batch effects”[39] caused by experimental procedure and human factors (exemplified with human saliva in this study). Most importantly, unlike traditional alignment-based method that highly depend on reference genome, QC-Blind could accurately identify and filter sequencing reads from target species utilizing only a small number of marker genes[40]. Moreover, the selection of marker genes is flexible and context-dependent, thus providing a lot of room for improvement during actual application.

As a future work, we are considering putting QC-Blind to the task where both target and contamination species are unidentified (and without biomarker genes other than 16S rRNA or a few genes) before sequencing. This problem is equivalent to the problem of metagenomic read binning, in which the reads of both target species and contaminations should be clustered in separate clusters. A more complex situation is with multiple species as target species, and multiple known/unknown contaminant species (but without multiple samples for comparison). Theoretically, contig binning can be directly applied to this multiple-species problem, and there are two points worth noticing: First, to increase the accuracy of marker gene identification, sequences from the same species have to be precisely clustered. Second, the performance of assembly and contig binning method should be stable among different situations, especially when they lack a dominant species. Then for filtration of contaminations, we could use database of samples as references to filter reads that could match (HMM-match) to the known contaminating samples, either based on fragment mapping or based on read similarity assessment.

## AVAILABILITY

Codes, manual and test data for QC-Blind are available for download at http://www.microbioinformatics.org/software/QC-blind/index.html.

## SUPPLEMENTARY DATA

### Supplementary File 1.zip

cluster_BS_1_5.xlsx; cluster_BS_1_10.xlsx; Cluster_BS+SC_1_10.xlsx; cluster_Ecoli_1_5.xlsx; cluster_Ecoli_1_10.xlsx; cluster_SA_1_5.xlsx; cluster_SA_1_10.xlsx; evaluation of BS_1_5.xlsx; evaluation of BS_1_10.xlsx; evaluation of BS+SC_1_10.xlsx; evaluation of Ecoli_1_5.xlsx; evaluation of Ecoli_1_10.xlsx; evaluation of SA_1_5.xlsx; evaluation of SA_1_10.xlsx

The clustering results and evaluations of simulated datasets of BS_1_5 (contaminated by HOB5), BS_1_10 (contaminated by HOB10), BS+SC_1_10 (contaminated by HOB10), Ecoli_1_5 (contaminated by HOB5), Ecoli_1_10 (contaminated by HOB10), SA_1_5 (contaminated by HOB5), SA_1_10 (contaminated by HOB10).

### Supplementary File 2.docx

Supplementary table 1| Identification Sensitivity of target and contamination species for QC-Blind-processed simulated datasets.

Supplementary table 2| Assembly Statistics for simulated datasets.

### Supplementary File 3.zip

Realdata_cluster_BS.xlsx

The clustering results of real datasets of AB_BS when cutoff of CONCOCT was set to 500bp. Realdata_evaluation of BS.xlsx

The evaluations of real datasets of AB_BS when cutoff of CONCOCT was set to 500bp.

## ACKNOWLEDGEMENT & FUNDING

This work is partially supported by National Science Foundation of China grant 31871334 and 31671374, Ministry of Science and Technology’s high-tech (863) grant 2014AA021502 and 2018YFC0910502.

